# Lilikoi: an R package for personalized pathway-based classification modeling using metabolomics data

**DOI:** 10.1101/283408

**Authors:** Fadhl M. Al-Akwaa, Sijia Huang, Lana X. Garmire

**Affiliations:** University of Hawaii Cancer Center, Department of Epidemiology, 701 Ilalo Street, Honolulu, HI USA 96813; Molecular Biology and Bioengineering Graduate Program, University of Hawaii at Monoa, Honolulu, HI, USA 96822

**Keywords:** metabolomics, pathway, classification, feature selection, machine learning, mapping

## Abstract

Lilikoi (Hawaiian word for passion fruit) is a new and comprehensive R package for personalized pathway based classification modelling, using metabolomics data. Four basic modules are presented as the backbone of the package: 1) Feature mapping module, which standardizes the metabolite names provided by users, and map them to pathways. 2) Dimension transformation module, which transforms the metabolomic profiles to personalized pathway-based profiles using pathway deregulation scores (PDS). 3) Feature selection module which helps to select the significant pathway features related to the disease phenotypes, and 4) Classification and prediction module which offers various machine-learning classification algorithms. The package is freely available under the GPLv3 license through the github repository at: https://github.com/lanagarmire/lilikoi

## Introduction

Metabolomics has been increasingly employed as a systematic approach to investigate the relationship between cellular signals and phenotypes [1]. Non-targeted metabolomics with global measurements, helps to discover novel metabolites biomarkers for diseases and conditions [2]. However, due to factors such as non-standardized protocols and highly heterogeneous study populations, it is difficult to find robust biomarkers that can be translated to clinical applications[3, 4].

Currently there are multitudes of secondary metabolomics analysis tools, primarily in the form of web tools. Very few comprehensive packages exist in R/Bioconductor, the dominant bioinformatics scripting language, in order to support metabolomics data analysis. Various modules of metabolomics pipelines have been implemented in other programming languages, including preprocessing [5], compound mapping [6], pathway networks [7], visualization [8], deep learning [9] and statistical enrichment analysis [10]. In particular, pathway based approaches have been applied widely in the metabolomics field. These methods use metabolites as biological entities to summarize to pathway-level statistics, and then calculate the overrepresentation of pathways compared to the background. However, none of these pathway-based methods entails personalized measurements for specific pathways. Moreover, these pathway-based methods are rarely integrated with classification algorithms for the purpose of metabolomics biomarker modeling.

To address the void above, we here introduce a new R package called Lilikoi (Hawaiian name for passion fruit), which specializes in personalized pathway measurement and classification prediction models. We present this tool in four modules: 1) Feature-pathway mapper, which standardize metabolite ID and map them to pathways. 2) Dimension transformation, which derive personalized pathway deregulation scores from metabolite profiles. 3) Feature selection, which provide the users with a range of feature selection algorithms to select significant features related to phenotypes. 4) Classification and prediction, which list a series of classification algorithms to derive machine-learning models and give predictions on testing data sets.

## Methods

### Overall design of Lilikoi

The Lilikoi package can be divided into four functional modules (Figure 1): Feature Mapper, Dimension Transformer, Feature Selector and Classification Predictor. In the first module, Lilikoi takes metabolites profiles data from the user as the input feature, and standardizes the metabolite names to various IDs in databases including KEGG, PubChem, HMDB and METLIN. After the mapping step, the second module transforms metabolite profiles to a comprehensive pathway deregulation score (PDS) matrix, based on the *Pathifier* algorithm [11]. The third module employs various feature selection algorithms to select key pathway *features* in the training set that are significantly related to phenotypes. The final classification module builds a classification model on the training set, based on various algorithms including random forest (RF), support vector machine (SVM), linear discriminate analysis (LDA), logistic regression (LOG), prediction analysis for microarray (PAM), generalized boosted model (GBM), recursive partitioning and regression analysis (RPART). It then performs prediction and quantitative evaluations on testing sets using various metrics. The details of each module are discussed in the following sections.

**Figure 1:**
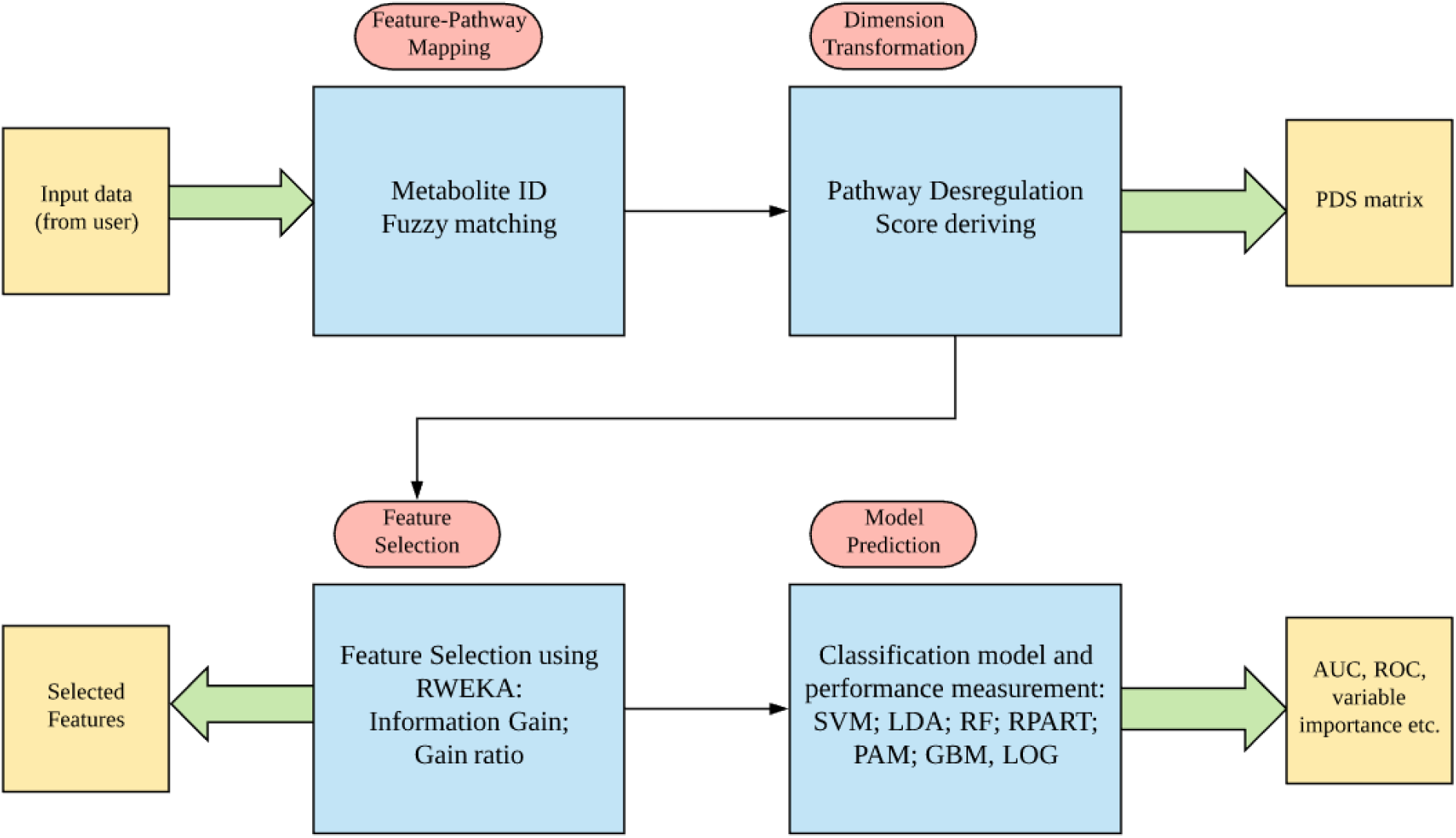
The workflow of Lilikoi package. Lilikoi composed of four modules: Feature Mapper; Dimension Transformer; Feature Selector; and Classification Predictor.

### Feature mapper

The feature mapping process consists three steps (Figure 2): In step 1, the input metabolite names are mapped to HMDB IDs using exact matching. We include various databases such as HMDB, KEGG, PubChem and MetaboAnalyst compound databases, to standardize the metabolite names. In step 2, Lilikoi employs the synonym database to standardize the rest of the unmapped metabolites to HMDB IDs. The remaining unmapped metabolites go through the third fuzzy matching step. We calculate the Levenshtein edit distance as a measurement of string similarity, and map the metabolite to the closest related standardized metabolites [12]. Such process allows for maximal mapping of input metabolites to standardized HMDB IDs.

**Figure. 2.**
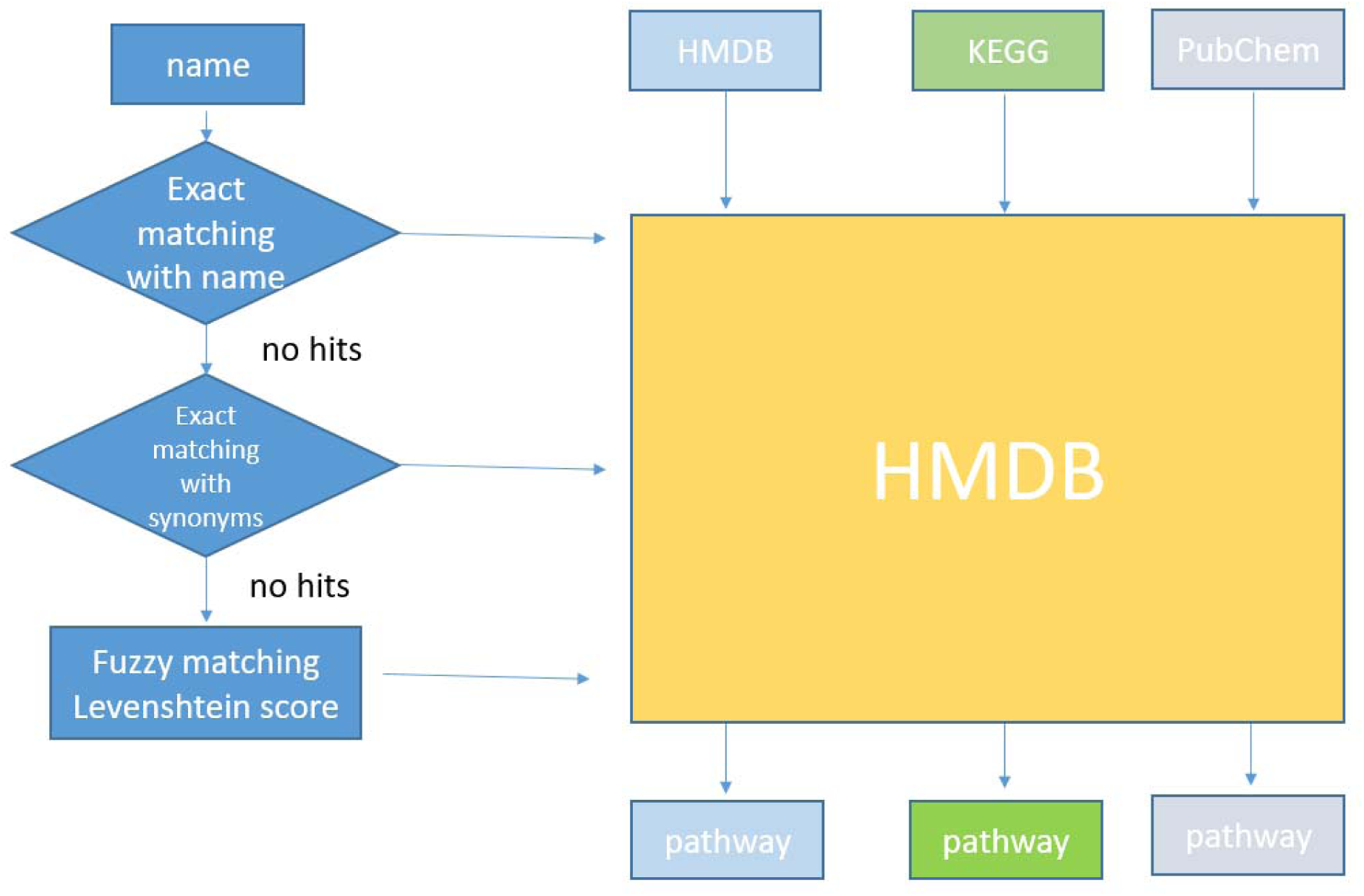
The workflow of Module 1: Feature mapper. User can input any metabolite IDs such as chemical name, KEGG, PubChem and HMDB IDs. Fuzzy matching algorithm is implemented to map the non-matched names to the 100k synonyms database.

### Dimension transformation

Lilikoi applies the *Pathifier* algorithm to perform the metabolites-pathway dimension transformation [11]. This algorithm summarizes per-sample information from the metabolite level to the pathway level [11]. For each pathway, all samples are mapped to a high dimensional principal component space (as data points) and a principal curve is constructed among them (the data cloud). A PDS score is then derived to measure the distance from the origin of the principle curve to the specific point on the principle curve, projected by the data point that represents a sample. The larger the PDS score is, the further a sample deviates from the normal level, in that specific pathway. As the result of the dimension transformation step, a new pathway-level metabolomics profile matrix is constructed. The users can then use this matrix for downstream analysis. More details of applications of *Pathifier* on biomarker studies (prognosis or diagnosis) can be found in our earlier publications[4, 13].

### Feature selection

Lilikoi allows the users to provide training and testing data sets, as well phenotype information for the samples. For the training set, Lilikoi provides two major feature selection algorithms: information gain (mutual information) and gain ratio, which select the most significant pathway-level features related to the phenotype. *RWeka* package is required for the feature selection module [14]. Information gain statistic is provided to evaluate the added information from each feature to help discriminate the phenotype. Gain ratio statistic is an alternative metric that solves the problem of overfitting, when there are a large number of distinct variables. We recommend the users use the gain ratio instead of the information gain when the input dataset has categorical variables besides the metabolomics data.

### Classification and prediction

Seven widely used machine learning algorithms, including LDA, SVM, RF, RPART, PAM, LOG and GBM are supported by Lilikoi, to build classification models. These methods have been widely used in the metabolomics community and reported in various research articles [9, 15–17]. Lilikoi uses R package *caret* for automatic parameter tuning of all the algorithms [18]. An n-fold (default n=10; flexible depending on different sample sizes) cross-validation is applied on the training dataset to avoid overfitting. Metrics to measure prediction accuracy, including Area Under the Curve (AUC), F1-statistic, balanced accuracy, sensitivity and specificity are reported to the user as barplots, similar to others [19]. Receiver operating characteristic (ROC) curves can also be reported as a separate figure.

### Combined model addressing confounding

Users can add any clinical factors such as age, sex and ethnicity to the model. All these factors are normalized between 0 and 1, by scaling between minimum and maximum values, so that they are compatible with the PDS score.

### Example dataset

For demonstration, we present a metabolomics data set from the City of Hope Hospital (COH) that was published earlier [4]. This dataset is composed of 207 samples from plasma (126 cases and 81 controls). The details of the data are summarized in our previous work [4].

### Package availability

The package is freely available under the GPLv3 license through the github repository at: https://github.com/lanagarmire/lilikoi

## Results

For illustration purposes, we applied Lilikoi to previously published breast cancer vs. normal control metabolomics data, which also have clinical information such as age, sex and ethnicity [4].

### Standardization and mapping of metabolomics IDs

We first used Lilikoi to transform the metabolite names to standard IDs. As different metabolomics research laboratories/preprocessing tools generate metabolomics profiles using different naming standards, Lilikoi allows the user to input any kind of metabolite IDs, their synonyms, KEGG IDs, HMDB IDs or PubChem IDs. Moreover, Lilikoi embeds comprehensive databases including over 18,000 metabolites and 100,000 synonyms, in accordance with other types of input IDs. Another major user-friendly characteristic of Lilikoi is the implementation of a fuzzy matching algorithm, which allows better mapping of uncertain metabolites, by calculating the string similarity score of the input metabolite name with those in the databases. These features of Lilikoi greatly improve its usability. In this example, 182 out of 227 metabolites are mapped to standard HMDB IDs.

### Metabolite to pathway level transformation

After transforming metabolites to standardized IDs, the metabolomics profile of the training set is transformed to a pathway-based profile through module 2: dimension transformation, with additional phenotype input (cancer/control) also provided by users. We the split the plasma data into 80% training and 20% hold-out testing set. In this example, the metabolites are mapped to a total of 93 pathways.

### Metabolomics feature selection

The next step is the feature selection module, using the PDS matrix and phenotypes of the training set as input. Users can choose either information gain or gain ratio to select key pathway attributes. Lilikoi plots a barplot of selected features and their relevance to phenotype labels. Lilikoi enables the output of information gain, a measure of feature relevance to phenotype for each selected attribute (Figure 3). In this example, nine pathways are identified as feature pathways in the plasma training set. Among them Alanine, aspartate and glutamate metabolism stands out as the most important pathway, with the highest information gain score.

**Figure 3.**
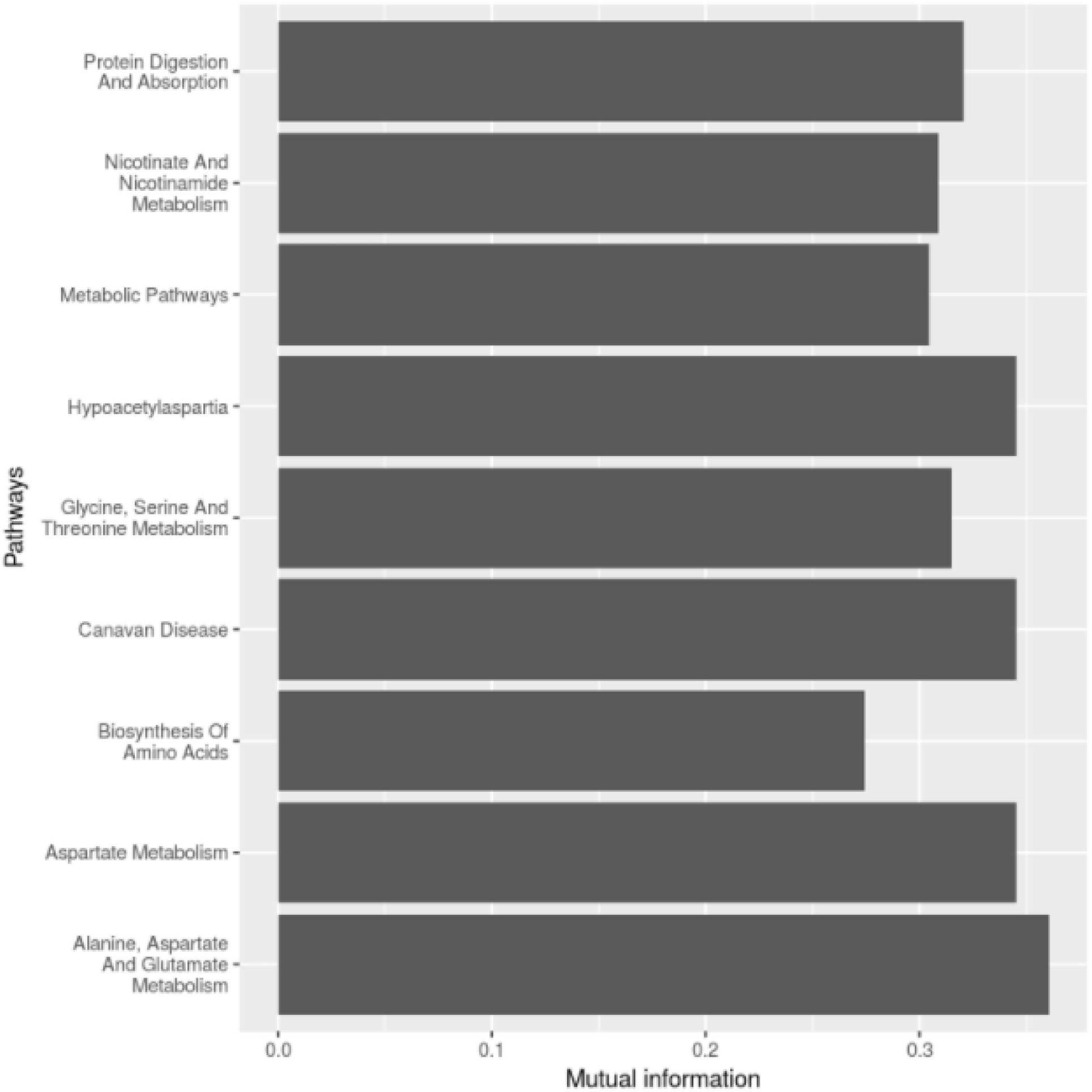
Plot of selected pathway features measured by Information Gain. The x-axis represents information gain score that measures the importance of the pathways, and y-axis displays the names of pathways selected from the training data.

### Model construction and validation

The last step is classification model construction and prediction. This module builds a model from the selected pathway features and allows the users to select among seven different classification algorithms with n-fold cross-validation. Users can compare performance measurements and choose the best classifier as the model of choice (Figure 4). This module generates two types of figures; a plot of ROC curves (Figure 4-A) which presents the overall model performance on the testing data set; and a second barplot (Figure 4-B) which illustrates the values of additional performance metrics (AUC, SEN, SPEC etc) of the testing data. In addition, Figure 4-C shows the performance metrics generated from the best-performed model, using a user selected the metric. In this example, we use AUC as the metric to select the best model, and GBM algorithm yields the best performance (Figure 4-C).

**Figure 4.**
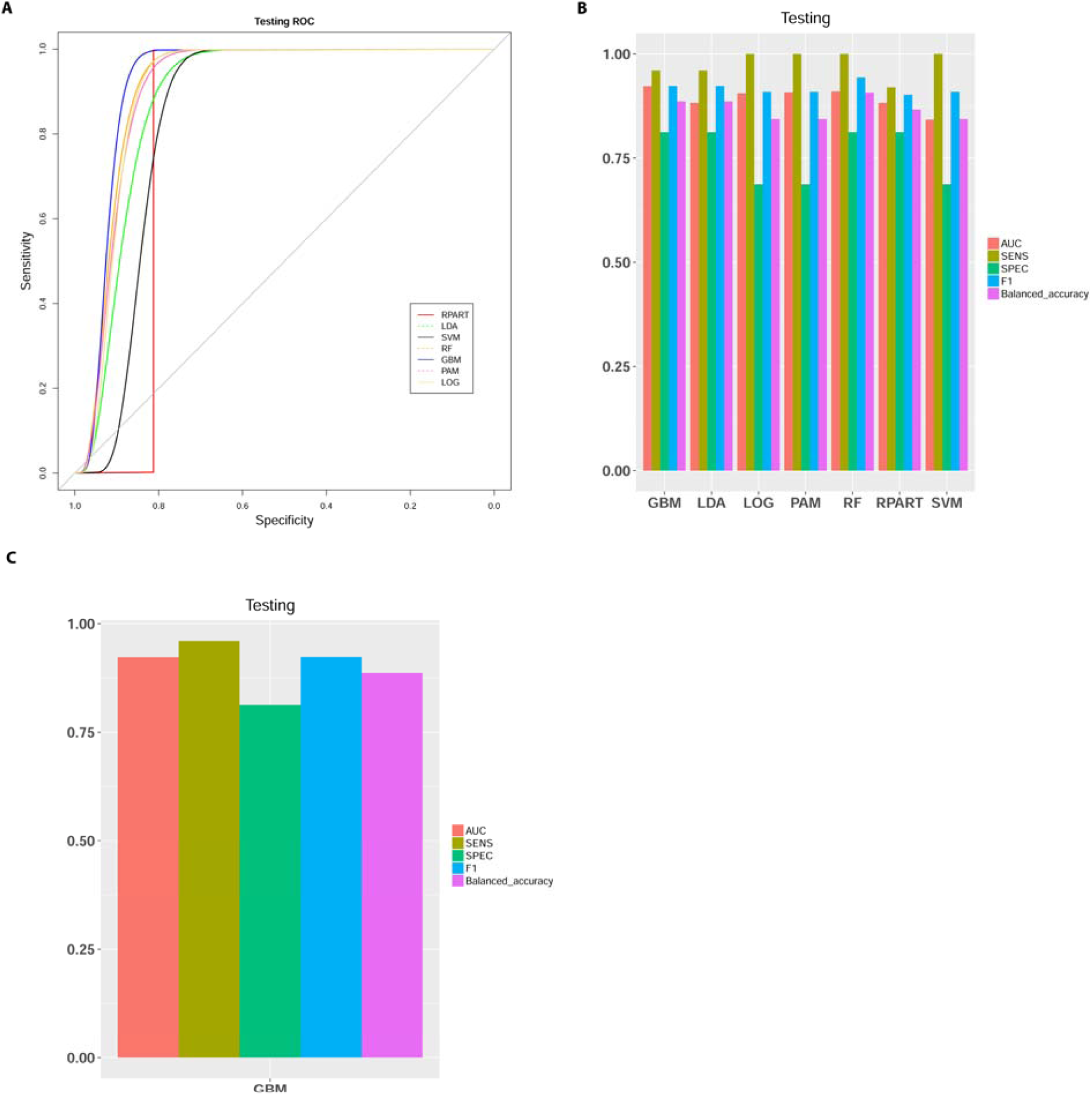
Model evaluation. (A) ROC curves of the breast cancer diagnosis testing set, obtained from seven classification algorithms: recursive partitioning and regression analysis (RPART), linear discriminate analysis (LDA), support vector machine (SVM), random forest (RF), generalized boosted model (GBM), prediction analysis for microarray (PAM), and logistic regression (LOG). (B) Metrics (AUC, MCC, Sensitivity, Specificity and F-1 Statistic) to measure the performance of classification on training or testing data? (C). Metrics of the best-performing model on testing data, based on the criteria chosen by user (AUC in this case).

### Model calibration by addressing confounding

Adjusting the fitted model using the clinical factors (if available) is a critical step in metabolomics based biomarker research. In this step, Lilikoi build three models on the training data set and plots the ROC curves on the testing set (Figure 5-A). Model 1 (black solid curve) is created using the selected pathways from the features selection module; model 2 (red dashed curve) uses the clinical factors selected by the user, and model 3 (blue solid curve) is created by combining both selected pathways and selected clinical factors. In this example, the clinical fators impose significant confouding in classification, and age is the primary contributor in the clinical model (data not shown). To understand the relationships among the selected pathways and the clinical factors, a correlation heatmap is plotted in Figure 5-B.

**Figure 5:**
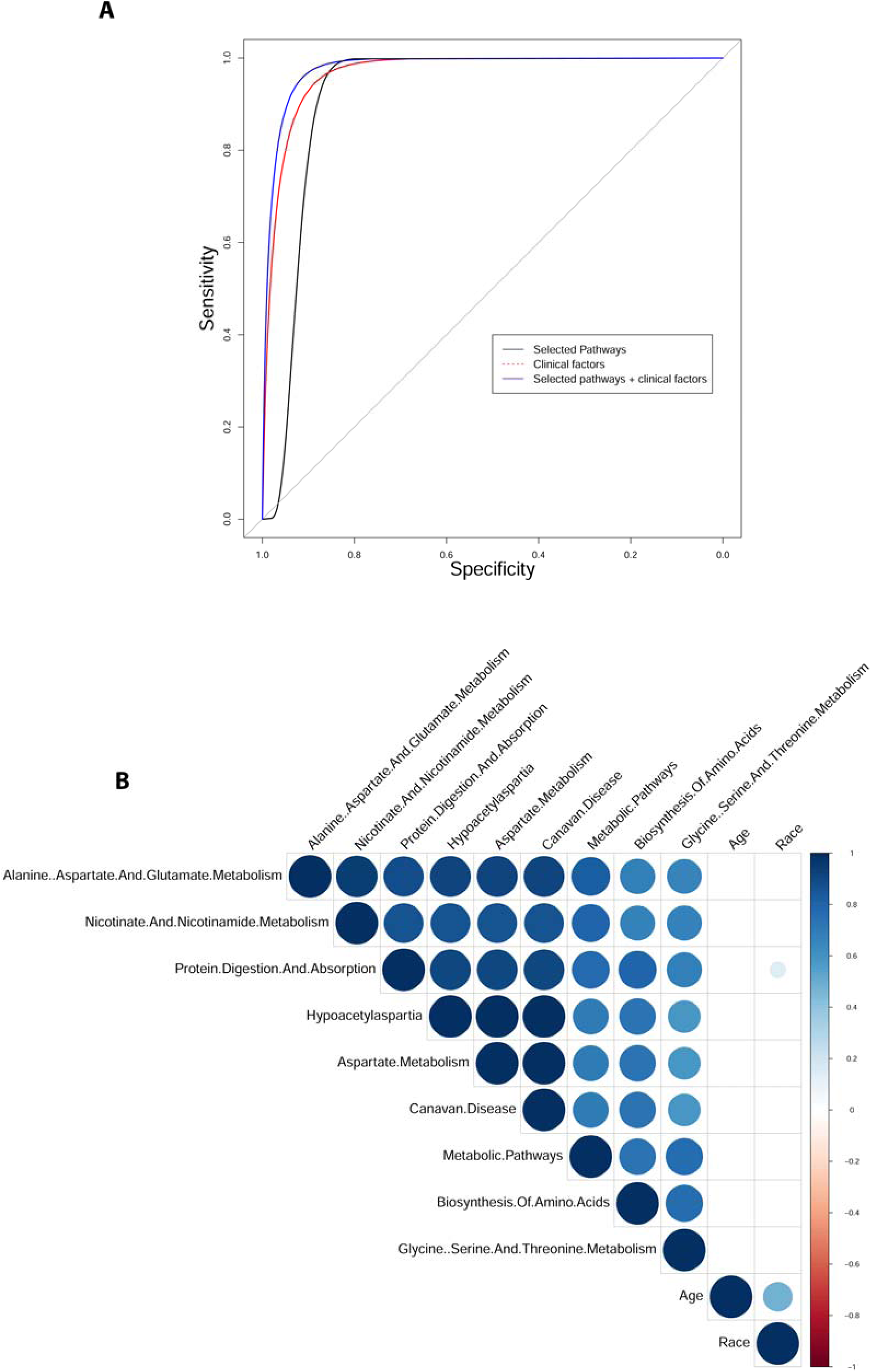
Calibration of metabolomics model by confounding. (A) ROC curves of metabolomics only, clinical data only, and metabolomics clinical combined model. (B) Correlation coefficients among demographical/physiological factors and the metabolomics data. Blue colors indicates positive correlations and red indicated negative correlations

## Discussion

Metabolomics biomarker discoveries have gained increasing amount of attention recently, in a variety of applications such as disease diagnosis and progression. Currently most of the biomarker features in metabolomics field are represented as individual metabolites, which suffer from inconsistency among studies. On the other hand, most pathway-based methods in the metabolomics field are not personalized and they are merely used for graphical mapping and enrichment analysis. None of these metabolomics pathway-based tools employ pathways as features for downstream biomarker modeling. Lilikoi addresses all these issues with personalized pathway deregulation measurements (PDS scores) and offers a standardized classification model for biomarker prediction. Compared to the traditional way of identifying individual metabolites as biomarkers, pathway-based biomarkers are more tolerant to population heterogeneity. Additional advantages of Lilikoi include the flexibility of its feature selection methods, the use of various machine learning classification algorithms, and its automatic tuning of parameters to generate the best model for a specific algorithm.

As an R package that will undergo actively improvements, Lilikoi can potentially benefit from other technical tweaking. Currently, a small percentage (20%) of the metabolites still cannot be mapped to the standard names in databases; also a fair amount of metabolites (40%) cannot be mapped to pathways, as the metabolite-pathway relationship is built upon pre-existing databases. One possible improvement could be using natural language processing (NLP) to improve metabolite ID standardization and metabolite-pathway mapping relationships, using the records in the literatures. Additionally, although the parameters in each classification model are automatically optimized, there is no automatic algorithm (AutoML) implemented that selects the best overall classification model; rather it depends on the user’s subjective preference of a machine learning method. It would be beneficial to automatically provide users references for classification algorithm selection, without human supervision [20, 21]. We plan to use AutoML in our classification module in the future.

## Consent for publications

All authors consent for publication

## Author Contributions

LXG envisioned the project, obtained funding, designed and supervised the project and data analysis. SJH and FMA implemented the package. FMA packaged the software, wrote documentations & instructions and generated Figures 2-5. SJH wrote the majority of the draft with help from FMA, and generated Figure 1. All authors have read, revised, and approved the manuscript.

## Acknowledgements

We thank all members in Garmire group for reviewing and commenting on the manuscript. We especially thank Xun Zhu for testing the package to ensure reproducibility of the results.

